# Socially learned arbitrary call use in a wild primate

**DOI:** 10.1101/2022.04.04.486940

**Authors:** Adwait Deshpande, Klaus Zuberbuhler

## Abstract

Human speech is an unusually flexible form of primate communication. A limited repertoire of phonemes is assembled into morphemes whose combinations are linked to referents by social learning in culturally specific ways. Non-human primate vocal utterances, in contrast, are drastically less flexible at every level, i.e., acoustic diversity, combinatoriality and referential function. Here, we were interested in whether primate vocal behaviour, despite its inflexible nature, is still amenable to social learning, a well-documented, powerful mechanism in many other aspects of primate behaviour. We provided wild vervet monkeys with social learning opportunities to operate a food dispensing apparatus by uttering specific vocalisations, ‘move-grunts’, that were contextually inappropriate. In two separate experiments, subjects witnessed how another group member produced move-grunts whilst operating a food dispenser. In experiment 1, subjects only heard playbacks of the calls, followed by food release; in experiment 2, they witnessed a complete demonstration video of conspecific including the calls, followed by food release. None of the subjects learned the task in experiment 1, but 1 of 39 subjects, a juvenile female, succeeded in experiment 2 by producing move-grunts at the food dispenser to obtain food rewards. We discuss our results and behavioural observations in relation to the current theory and its eventual implications for reconstructing the evolutionary pathway to human speech.

## Introduction

Human language is the capacity to represent and communicate events as hierarchically structured, recursively nested mental structures via a linear signal stream [1]. The natural vehicle for this process is speech, which consists of language-specific phoneme repertoires that range from less than a dozen (Piraha) to over a hundred (⍰Khong). Each language can be phonetically transcribed in terms of its speech sounds [2] and in how sound combinations contribute to linguistic structures, making human speech arguably the most flexible and complex natural vocal communication system known [3,4]. The underlying ability to generate speech sounds emerges early during human ontogeny, around the end of the first month, with infants producing proto-phones decoupled from any affective state and basic biological function, the ‘babbling’ behaviour [3,5]. Proto-phones are produced in addition to a more standard repertoire of functionally fixed vocalisations such as laughing, crying or protesting [6], used in specific social interactions similar to the calls of non-human primates. Proto-phones, however, do not have any obvious biological function but appear to be the foundation for subsequent speech development. During this process, infants socially learn from other group members to use these sounds and their combinations to interact socially and as referents for external events [7,8]. Thus, to reconstruct the evolutionary history of speech and language, it is vital to shed light on how human evolved the capacity to produce functionally flexible sounds amenable to social learning.

One promising way to address evolutionary questions of human behaviour is to make direct comparisons with closely related non-human primates [9]. According to most theories, primate communication is characterised by small call repertoires, with signallers prevented from majorly modifying the acoustic structure of existing calls and from producing novel sounds [10,11]. Moreover, apart from reports on marmosets (Reviewed in [12]), non-human primate infants do not show human-like babbling behaviour but gradually acquire an adult-like vocal repertoire with no compelling evidence of social learning [13]. Flexibility is somewhat higher in primate call usage [14], with examples of signallers using the same call in different contexts. For example, bonobos (*Pan paniscus*) produce ‘peep’ calls across several contexts, encompassing various emotional valences [15]. The same has been found for the ‘grunt’ calls in juvenile chimpanzees (*Pan troglodytes*) [16] and ‘coo’ calls in different macaque species [17–19].

While these findings confirm that primates can use the same call in different contexts, it is still possible that call production is the direct result of the same underlying psychological state experienced by callers. For example, many primates produce acoustically distinct alarms to a wide range of terrestrial disturbances, including non-predatory events [20], suggesting the same basic inner state evokes the calls [11,21]. However, other primate calls, such as bonobo ‘peeps’ or macaque ‘coo’ calls, are given to such a wide range of events that they appear to be affectively decoupled, i.e., functionally flexible [15,17,18]. In an early study, Sutton et al. were indeed able to condition rhesus macaques (*Macaca mulatta*) to produce ‘coo’ calls following arbitrary visual stimuli [22], although the test window was very long (5 min after stimulus presentation), suggesting that coo calls might have been produced for other reasons (see [22,23] for similar concerns). In other studies, conditioning has been possible with food calls. However, here, the issue is that callers may produce the calls in anticipation of the food rewards when completing a trial successfully, not as a response to the conditioned stimuli [17,24]. A recent study with rhesus macaques attempted to overcome these limitations by showing that subjects could switch between two call types based on different visual stimuli [25].

While such experiments demonstrate that non-human primates have some degree of volitional control over call use, they all required extensive training in captive settings with operant conditioning paradigms, raising questions about ecological validity. In natural conditions, the conditions for operant conditioning are rarely met, as both social and physical environments change continuously while consistent reward patterns are unlikely to be found. Social learning, i.e., learning by observing others, is a much more probable learning mechanism, which is well-documented and highly adaptive in wild animals, especially in foraging or antipredator contexts [26,27]. However, its importance in call use and communication is not well documented [28].

In this study, we addressed this issue by providing social learning opportunities to wild South African vervet monkeys (*Chlorocebus pygerythrus pygerythrus*), a species with an established capacity for social learning [29–31] and a rich vocal repertoire of calls with potential for functional flexibility [32,33]. We conducted two experiments on two different groups of monkeys that introduced subjects to the fact that a food dispenser could be activated by producing an out-of-context vocalisation, a ‘move-grunt’. In the experiments, subjects witnessed another group member obtaining food from the dispenser by producing move-grunts (or ‘moving-into-an-open-area’ grunt), a call type typically given by individuals that move into open areas, observe other group members do so [34] or initiate group movements (AD, unpublished data). The call emerges relatively early during ontogeny, with 8-12 weeks old juveniles already producing the call in the correct context [35]. Due to these observations and the fact that this call type does not provoke any visible positive or negative affect in callers or receivers [36], we speculated that vervet monkey ‘move-grunts’ could be functionally flexible call, similar to bonobo ‘peep’ calls or macaque ‘coo’ calls. And possibly more amenable to social learning than other vocalisations.

In the first experiment (conducted on the first group of monkeys), subjects heard a playback of a move-grunt by a familiar group member directly followed by a food reward delivered from a food dispenser (Supplementary material Figure S2). We predicted that if monkeys were able to learn the association between call type and food reward, they would start producing move-grunt themselves to get the rewards. In the second experiment (conducted on the second group of monkeys), subjects watched a demonstration video of a conspecific obtaining a food reward following playback of a move grunt. This second experiment is based on findings in both wild and captive settings, showing that non-human primates can learn to extract food from an apparatus [37,38] and that they can learn from watching video clips. Again, we predicted that subjects could learn to operate the food dispenser from observing others producing move-grunts.

## Methods

### Study site and groups

The study was conducted at the Inkawu Vervet Project (IVP) on the Mawana Game Reserve, Kwazulu Natal, South Africa (S 28° 00.327; E 031° 12.348). At the IVP, several neighbouring groups of wild South African vervet monkeys (*Chlorocebus pygerythrus pygerythrus*) are well habituated to human presence and allow close observations by multiple observers. All observers go through standard training followed by inter-observer reliability tests for individual recognition and ethogram recordings before participating in data collection. Individuals were fed regularly on corn kernels and apple pieces for various experiments. Food provisioning events were announced using specific calls by observers, which are understood by monkeys [27,39].

### Experiment 1: Social learning of auditory demonstrations

In the first experiment, we provided monkeys with a playback call of another group member’s move-grunts immediately followed by a food reward delivered manually from a dispenser. This experiment was conducted on group Noha (NH). At the time of the experiment, the group’s size was 34 individuals, including juveniles. We used a naturally recorded move-grunt of a middle-ranking adult female of the group as a playback stimulus. The call was recorded when the female was moving out from the sleeping site in the morning (see Appendix). To record the call, we used an audio recorder (Marantz PMD 661) and a directional microphone (Sennheiser MKH 416 P48) with a sampling frequency of 44.1 kHz. We padded the call with 2 s of silence before and after using the Raven Pro 1.5 for the playback [40].

For the delivery of rewards, we used a custom-made, manually operated food dispenser. It was made of a handheld (1.5 m long and 7.6 cm diameter) polyvinyl chloride (PVC) pipe and a translucent plastic container. The experimenter sat on a field chair holding pipe at one end while the other end of the pipe was positioned inside the plastic container 1-2 m away from the experimenter (Figure 1). The experimenter was holding hidden food rewards (red apple pieces approx. 2.5 cm cubes). To reward a subject, the experimenter dropped an apple piece from his/her end of the pipe. The apple piece slid down the pipe and came out from another end in the plastic container to become accessible for the subject.

**Figure 1:**
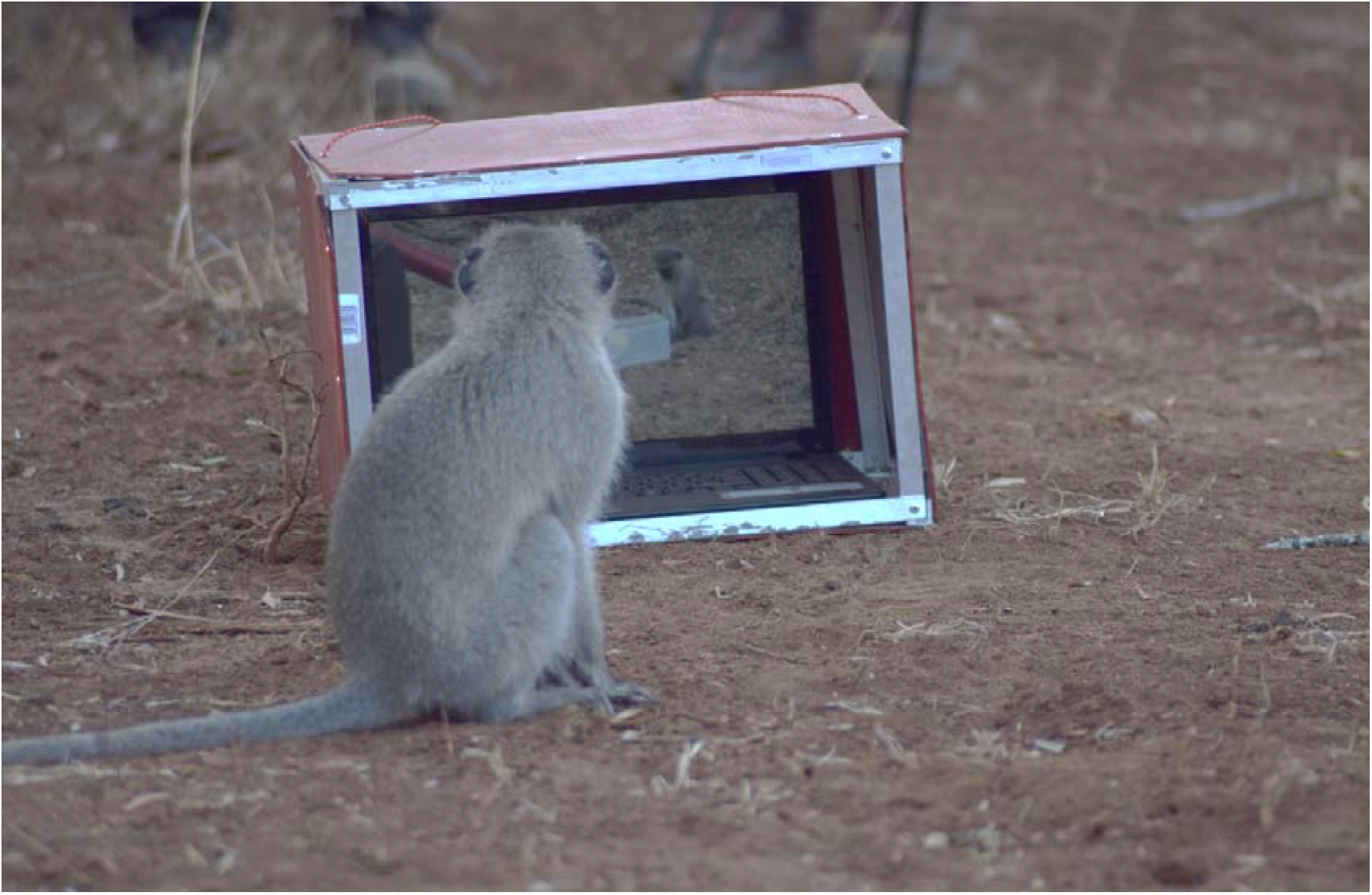
Photograph of a subject monkey watching a demonstration video during Experiment 2: Social learning of audio-visual demonstrations

The final experimental setup consisted of the food dispenser handled by experimenter A1 (see Appendix). The hidden speaker (Nagra Kudelski, DSM-Monitor) was placed 5m away from the food dispenser, and the playback call was played from a speaker through a Motorola Xplay phone. We pre-adjusted the speaker amplitude similar to a natural call which was kept constant throughout the experiment. The speaker was operated by experimenter B1 sitting close to it. Experimenter C1 who recorded the subjects’ behaviours, was positioned 3-8m from the food dispenser. We used two cameras (JVC quadproof EverioR camcorder) on tripods to record all the experimental sessions. One camera was positioned to cover the entire setup with a wide-angle view, whereas the other camera was focused on the food dispenser. We also recorded all the vocalisations elicited around the food dispenser using a solid-state audio recorder (Marantz PMD 661) and a directional microphone (Sennheiser MKH 416 P48) with a sampling frequency of 44.1 kHz.

Each experimental session started early in the morning near the sleeping site of the monkeys just after sunrise. Once the experimental setup was ready, we attracted the monkeys to the setup using conventionalised human ‘food-provisioning calls’ (see study site and groups) and provided them with a small number of corn kernels for the first three sessions. Once more than one monkey was present in the setup’s proximity (5-10m), we started an experimental session. We conducted seven experimental sessions, which lasted from 10 to 105 minutes (Mean 56.8 minutes). A session ended when the group moved away from the setup.

Each experimental session consisted of training and contrast trials. For the training trials, experimenter B1 played a move-grunt from the hidden speaker in response to which the experimenter A1 immediately (< 1s) dispensed a reward usually picked up by the subject. If no individual approached the dispenser, the rewards were recovered by experimenter A1; such trials were considered trials without subjects. We gradually increased the time between two consecutive trials with each experimental session (mean: 54s, range: 1s-698s). Food rewards were also dispensed when a monkey in the proximity of the setup (5-8m) naturally elicited a move-grunt.

For the contrast trials, the experimenter dispensed non-edible items (small stones) from the dispenser without a playback. Contrast trials were designed to ensure that monkeys did not associate subtle, unintentional cues from the experimenter with the appearance of food rewards, i.e., to prevent Clever-Hans effects [41]. Contrast trials were conducted between two training trials randomly decided by the experimenter A1.

For each trial, the monkey closest to the food dispenser was considered the subject. If multiple monkeys were present, the experimenter considered the first individual looking at the dispenser as the subject.

For each trial, the experimenter C1 recorded the following behaviours by the subject: (1) Look at the speaker (yes/no: head orientation towards speaker after playback); (2) Look at the dispenser (yes/no: head orientation towards food dispenser before the reward was dispensed); (3) Gaze alternation (yes/no: head orientation towards speaker followed by rapid (<2s) reorientation to the dispenser (or vice-versa; see supplementary <u>Video</u>); (4) Startle response (yes/no: abrupt, sudden and brief locomotor response. For each trial, the second observer noted the identities of the audience members within a 5-8m radius around the setup.

We considered social learning to be accomplished if a subject elicited a move-grunt in the proximity of the food dispenser with a head orientation towards the dispenser and without any apparent group movement immediately before and after the call.

### Experiment 2: Social learning of audio-visual demonstrations

In the second experiment, we provided monkeys with an opportunity to learn the novel usage of move-grunt from a demonstration video. This experiment was conducted on the ‘Baie Dankie’ group (BD) with 64 individuals, including juveniles. The demonstration video was edited from footage of two different events from Experiment 1. In the videos, a subject monkey from group NH can be seen stationed at the food dispenser and rewarded with an apple piece after the playback of move-grunt. We then removed the original audio track from both the footages and manually inserted the move-grunt (same as playback of Experiment 1) just before the food reward was dispensed in both the footages. Editing made both video events appear like the stationed monkey is eliciting the move-grunt and getting rewarded immediately after the call (at least to the human observer). In both the footages, the experimenter providing the food rewards was only partially visible. We then stitched one footage after another in a loop to make a 30 min long demonstration video showing 120 demonstrations (60 demonstrations of each event) (see Supplementary <u>Video</u>). The demonstration video was played on a 15-inch LCD laptop screen. The laptop was placed inside a protective opaque box with transparent plexiglass on one side for video watching (henceforth a video box).

The final experimental setup contained two components; the video box place on the ground and the food dispenser (same as Experiment 1) held by the experimenter A2. The food dispenser was placed in front of the video box, approximately 3-5 m away. We also placed two different video cameras (JVC quadproof EverioR camcorder) on tripods. One camera recorded the whole setup with a wide-angle view. The second camera, along with the audio recorder (Marantz PMD 661) and a directional microphone (Sennheiser MKH 416 P48), was placed to record the behaviours and vocalisations of the monkeys around the food box (see Appendix). Each experimental session was 45 mins. The video was played for 30 mins, and for the last 15 min, we waited at the setup without video playing. Monkeys were free to interact with the food dispenser throughout the entire session. We did 2-5 playbacks of the move-grunt from a hidden speaker (Same as Experiment 1) in the last fifteen minutes of the seven experimental sessions. The experimenter dispensed the reward after each playback which was picked up by a subject monkey. The objective was to reinforce that the reward is independent of the videos and contingent solely on call elicitation. Two other experimenters B2 and C2, were present during each experimental session. Each experimental session was conducted in the early morning near the sleeping site of the monkeys. We chose a relatively open space near the sleeping site to set up the experiment. Before starting the videos, we attracted the monkeys to the video box by putting a few corn kernels in front of the box. The experimenter A2 played the video on the laptop and assumed the food dispenser position, which marked the start of the experimental session. During eight sessions, the monkeys left the area within 45 mins. We waited for 10-15 min for the monkeys to come back to the area; otherwise abandoned the session before 45 min. We conducted one experimental session every week over three months, with a total of 20 experimental sessions.

During the session, the experimenter A2 rewarded a subject upon the successfully eliciting move-grunt. I.e. if the subject elicited a move-grunt in the proximity of the food dispenser with a head orientation towards the dispenser and without any apparent group movement immediately before and after the call (see supplementary <u>Video</u>). The experimenter was highly trained in identifying call types in this species; however, as required to dispense the reward immediately after the call, there was a margin for human error (See results).

We also recorded the following behaviours during each experiment session. (1) Video-watching: Sitting/standing directly in front of the video box with the direct, clear head orientation towards the laptop screen when the video was playing. We recorded all the individuals looking at the video during the scan taken every minute by experimenter B2 [42]; (2) Look at dispenser: Direct gaze which inferred through a clear head orientation towards food dispenser while video-watching. The dispenser was placed opposite the video box, making it easier to infer clear head orientation as the subject had to turn around the head more than 90 degrees. (3) Approach the dispenser: The monkey was considered approaching the dispenser when it walked directly towards the dispenser. The last two behaviours were recorded ad-libitum by experimenter C2. Any monkeys that elicited one of the three behaviours were considered subjects for the analyses. We used a long-term dataset of the BD group’s vocalisations to determine the natural frequency of move-grunt calls. These call recordings were collected over 36 days spread across six months, with a sampling effort of 8 hrs per day. We recorded all the vocalisations elicited by BD group members during the sampling, which the observer could hear. For both experiments, descriptive statistical analyses were conducted in R v 4.0.3 [43].

## Results

### Experiment 1: Social learning of auditory demonstrations

We carried out a total of 440 trials (392 training, 48 contrast trials) distributed over seven experimental sessions (training: median= 56, mean = 62.8 ± 17.92, range 11-133 trials per session; contrast: median= 6.5, mean = 8.0 ± 2.73, range = 2-22 trials per session) between sessions two and seven. In 344 trials a subject was present (range 3-11; median= 4, mean= 5.57 ± 1.28 subject per session); the remaining 96 trials were without subjects. A total of 13 subjects participated in one or several trials (range 1-79, median= 15 trials, mean= 26.46 ± 1.64 trials per subject) (Table 1).

**Table 1:** Summary statistics for Experiment 1: Social learning of auditory demonstrations

| Total number of trials in seven sessions | Trials with subjects/<br>trials without subject | Total number of subjects | Number of times task was successfully completed |
| --- | --- | --- | --- |
| 440<br>(392 training trials,<br>mean= $62.8 \pm 17.92$<br>trials per session and<br>48 contrast trials,<br>mean= $8.0 \pm 2.73$ ) | 344/96 | 13<br>(mean= $5.57 \pm 1.28$ subject<br>per session) | 0 |

No subject completed the task successfully, i.e., no one produced a move-grunt between the two trials in any session while stationed at the dispenser with a direct head orientation towards the dispenser.

The participation by subjects was highly skewed. Three subjects participated in 229 trials (66.5% of trials). The first subject, Gran, was a high-ranking juvenile (4-year-old) female. The second subject, Wolf, was an adult male, whereas the third subject, Ula, was a juvenile (3-year-old) male. Due to this participation skew, we focused our subsequent behavioural analyses only on these three subjects. (See Appendix). We observed some trends in our raw data during the first session in terms of looking at the dispenser, looking at the speaker, no reaction and a startle response. From the second session, we never observed a startle response and looking at the speaker. The quick sequence of looking at the box and speaker or vice-a-versa was observed in session two, three and four. We only observed looking at the box or no reactions from the subjects in the last three sessions (Figure 2).

**Figure 2:**
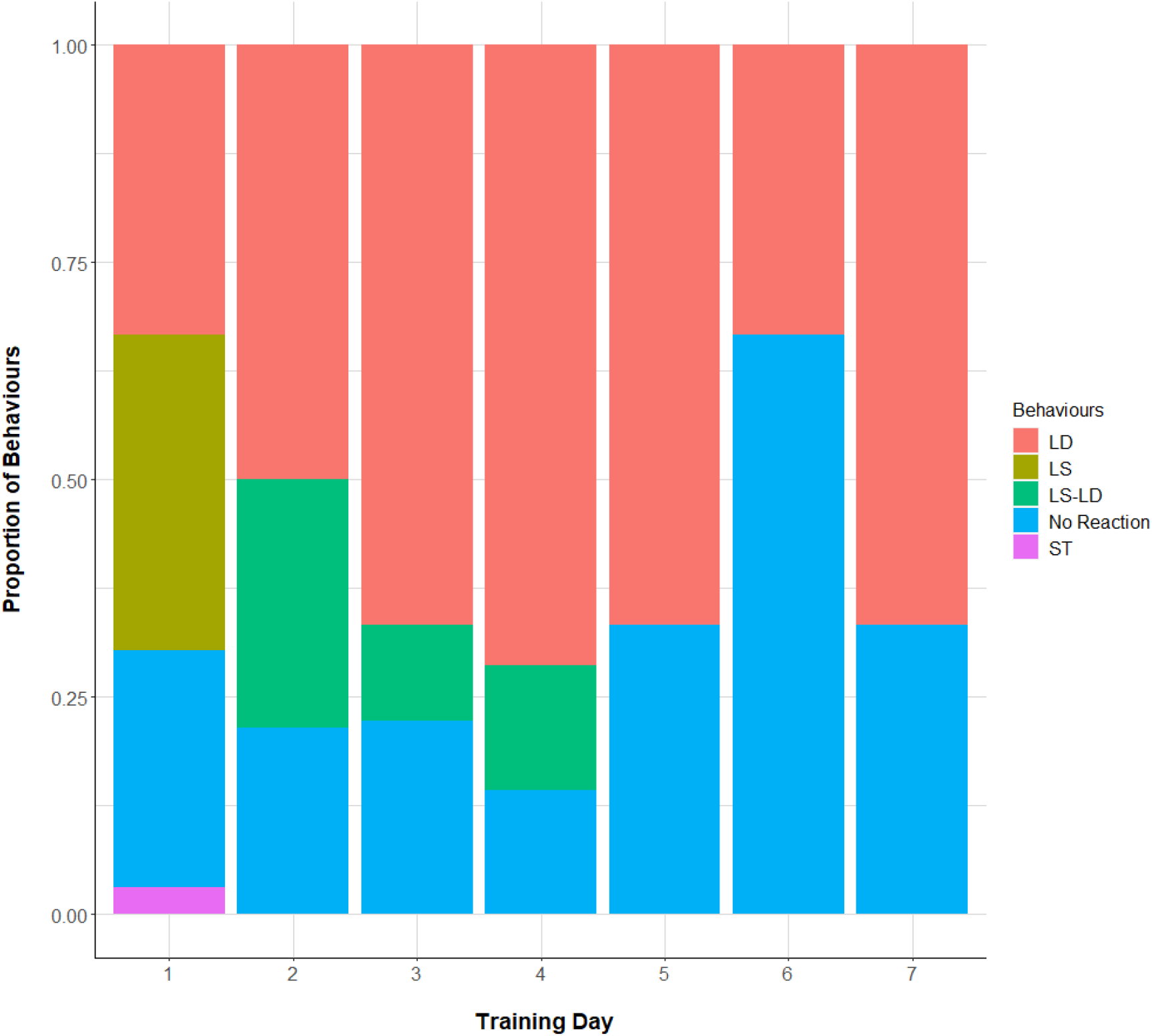
Proportions of cumulative behaviours of three subjects during participated trials across sessions of Experiment 1: Social learning of auditory demonstrations. LD (look at dispenser), LS (look at speaker), LS-LD (Gaze alternation), ST (Startle response)

### Experiment 2: Social learning of audio-visual demonstrations

We carried out a total of 20 sessions, which lasted from 20-57 min (Mean = 39m 51s ±2m). Ten experimental sessions were 45 min long as per the experimental design. The number of subjects per session ranged from 3 – 27 (median= 9, mean = 9.70 ±1.29 subjects per session). We analysed three behaviours elicited by the subjects in every session. Video-watching ranged from 4 – 138 times per session (median= 9.50, mean = 18.25 ±6.53 times per session). Looks at the dispenser ranged from 1-60 per session (median= 4.50, mean = 11.95 ±4.09 times per session). Approaches to the dispenser per session ranged from 1-34 (median= 10, mean= 11.35 ±1.77 times per session). Total behaviours (sum of the three behaviours) per session ranged from 11-197 (median= 25.5, mean= 42.1 ±10.20 behaviours per session) (See supplementary behaviour). Thirty-nine individuals participated in the experiment (adults: females N=9; males N=9; juveniles: females N=11; males N=10) (Table 2).

**Table 2:** Summary statistics for Experiment 2: Social learning of audio-visual demonstrations

| Total number of sessions | Mean duration of session ± Standard error | Total number of subjects | Total number of subjects who successfully completed the task | Total number of times task was completed successfully |
| --- | --- | --- | --- | --- |
| 20 | Mean = 39m 51s ±2m | 39<br>(mean = 9.70 ±1.29 subjects per session) | 1 | 33<br>(All the successful completions between session 12 to 19) |

One subject, Oorties (OT), a 4-year-old juvenile female, completed the task successfully by eliciting move-grunts in the previously specified way (see Appendix). A successful performance was observed from session 12 to 19, during which she produced 42 successful calls when stationed at the dispenser, for which she was rewarded every time by experimenter A2. Thirty-three calls were move-grunts produced in the correct way (Mean 4.12 correct calls per session between session 12 to 19). Further, three calls were also move-grunts but produced while OT was looking away from the speaker. The remaining six calls were possibly eagle alarm calls (for which she was wrongly rewarded by the experimenter; see supplementary <u>Video</u>).

OT watched the video 56 times across all sessions, which was the highest count compared to the other subjects (Figure 3). There was a significant Spearman’s correlation between cumulative successful calling and video-watching by OT (r = 0.91, p<0.001; Figure 4). In 10 cases, OT did not produce any audible vocalisations, but her body posture was as if she were to produce move-grunts (sitting and bending body forward towards the ground) with noticeable mouth and lip movements (see supplementary <u>Video</u>). These failed attempts were only observed during sessions 12 to 19, where she also produced at least one successful call. During 288 hrs of the sampling effort before this experiment, we recorded 189 natural vocalisations, out of which 17 were move-grunt calls. The natural frequency of move-grunt for a BD group was 0.05 move-grunt/hr.

**Figure 3:**
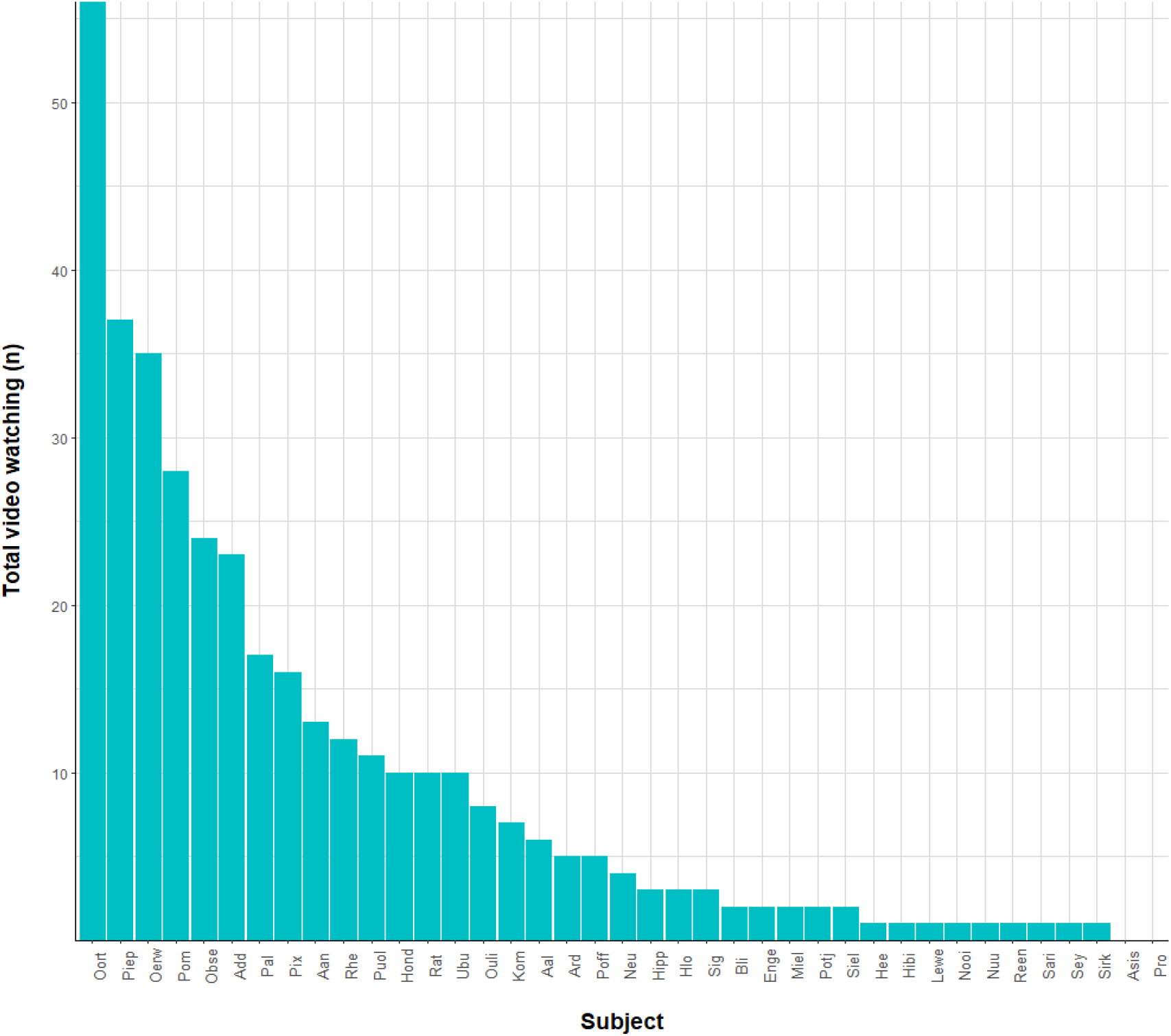
Total number of times subjects watched the video in all the experimental session of Experiment 2: Social learning of audio-visual demonstrations

**Figure 4:**
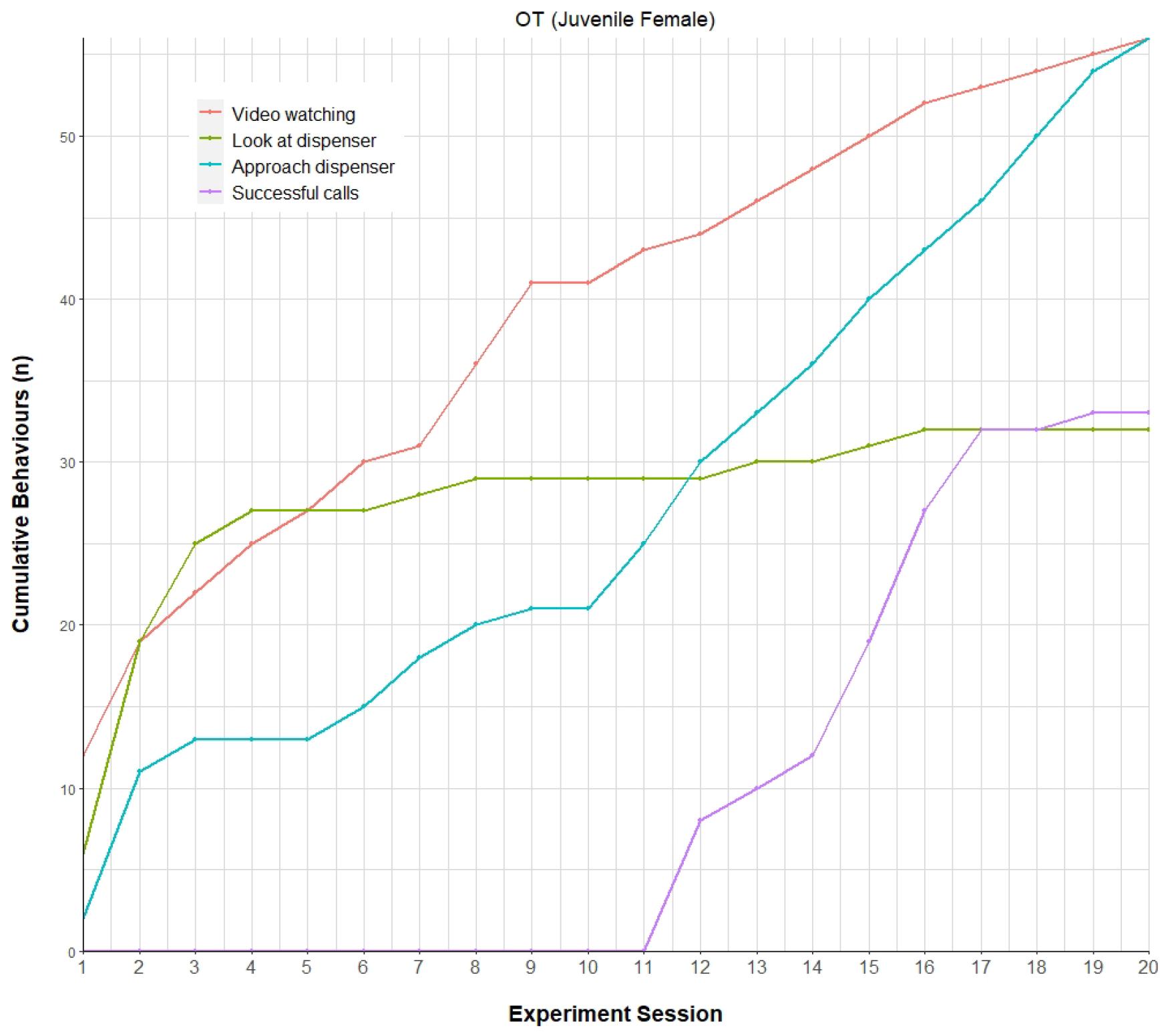
Cumulative behaviours by subject OT in Experiment 2: Social learning of audio-visual demonstrations

## Discussion

Human language is the product of human cognition, partially shared with animals, while speech appears to be uniquely human. No other species has the capacity to fluently combine phonemes into morphemes into phrases and sentences to not only interact socially but also to refer to real and imagined events [1]. This ultra-flexible sound production system is possible because of decoupling of vocal behaviour from direct biological function, insofar as humans are able to not only produce sounds that are linked to specific psychological states, such as distress is linked to crying, but additionally have the ability to produce sounds voluntarily [5,7]. Unhinged from biological function, human speech sounds are highly susceptible to social learning, which ultimately enables humans to establish countless signal-referent relations and connect with their minds.

This study attempted to explore the possible evolutionary roots of functionally decoupled and psychologically flexible calls in the primate lineage. To this end, we investigated if wild vervet monkeys can socially learn to produce affectively neutral move-grunt calls in an entirely novel context, i.e., to imperatively operate a food dispenser in their favour. To test this empirically, we provided wild vervet monkeys with opportunities to socially learn how to operate a food-dispenser, i.e., by producing spatially oriented move-grunts in order to activate the mechanism and receive a high-value food reward.

In a first experiment, we only provided a fragmentary glance of the event’s causal structure, i.e., by playing another group member’s move-grunts which was immediately followed by the release of food from the dispenser. Even though the event contained the key components of the causal chain, we did not find any evidence of learning in subjects. We concluded that a mere stimulus-response contingency was insufficient for subjects to adapt their own behaviour and successfully complete the task. In the second experiment, we featured the entire event, i.e. by showing a video clip of a monkey performing the entire event. Here we found that one monkey learned the contingency and spontaneously started to produce move-grunts in this novel condition, probably induced by watching the demonstration video.

In the first experiment, we played move-grunts from a hidden speaker followed by an immediate food reward from the dispenser. We expected that subjects would learn the association between move-grunt and food rewards and eventually start to produce move-grunt themselves to obtain the rewards. However, none of the subjects produced any move-grunt when stationed at the dispenser, suggesting that observing the stimulus-response contingencies was insufficient to learn a new call usage. Nevertheless, in three subjects that participated well, we observed behavioural pattern that indicated that some learning had taken place. On the first training day, monkeys were generally startled by the playback calls and looked at the speaker or the dispenser. On the following three days, however, the three subjects rapidly looked at the speaker first and then at the food-dispenser or the other way around. On the last three training days, subjects only looked at the dispenser and no longer at the speaker, indicating that the subjects learned the association between playback and subsequent food reward. Rapid auditory learning of call-event pairing has been found before in different primate and other mammal species [44–47].

In the second experiment, the subjects were able to watch a demonstration video of a conspecific successfully retrieving food by producing move grunts. The setup in this experiment was identical to the food dispenser nearby with which monkeys could interact freely. Out of 39 subjects who watched the video, one 4-year-old juvenile female (OT) started to produce move-grunt calls when stationed at the dispenser, a total of 33 times from session 12 onwards. Interestingly, she watched the video substantially more often than any other subject who participated in the experiment, suggesting that OT learned to produce a call in an arbitrary context by watching another individual doing the same, evidence for primary level usage learning [48].

An argument against this interpretation is that OT produced the first move-grunt for reasons unrelated to the video experience (e.g. by observing another individual moving at the same time) but then became conditioned due to the provisioning, learning from the video. However, this explanation is unlikely due to the fact that standard operant conditioning paradigms usually require enormous numbers of trials for subjects to become conditioned [22,23,25]. Furthermore, the natural rate of move-grunt producing is extremely low (1 move-grunt per 20 hours of observation per group), which makes it highly unlikely that OT produced her move-grunts due to a naturally adequate context. Finally, we did not observe any group movement during or after any successful move-grunts produced by OT. Also, OT did not move away with any group member immediately after any successful call. Thus, we conclude that our interpretation that OT learned a novel usage of move-grunts is much more plausible than alternative explanations. It is possible; however, that consistent reinforcement after move-grunt production strengthened the establishment of this new behaviour, especially in the beginning. In conclusion, this is the first experimental study of a socially learned novel call use in wild primates.

This finding is important because while earlier studies show that monkey vocalisations can be conditioned in controlled laboratory conditions [22,49–51], the operant conditioning paradigm used in such studies is not pertinent to the natural conditions in which primates live. We show that an alternative mechanism of observational social learning is likely to play an essential part in developing communicative competence [52]. Historically, social learning in primates has mainly been investigated in the acquisition of manual skills usually related to food acquisition [53,54]. But primates constantly monitor others during social interactions [55,56], suggesting that social learning could also play a role in becoming competent communicators. Future fine-grained studies on ontogeny and the interplay between social cognition and communication are required to test this possibility [57]. Social learning theory may have to be broadened to include all types of behaviour and communication [58–60].

Although the sample size is not optimal for comparing the outcomes of the two experiments, it is important to speculate why the demonstration video was probably more effective as a social learning stimulus than the playback of the call? Although the playbacks reliably predicted the availability of food, subjects were unable to see the causal connection, i.e., a conspecific uttering the call to trigger the production of food. Thus, the playback condition functions similarly to the ‘ghost control’ in manual tasks, with an invisible model, which may be cognitively more demanding for subjects [61]. The importance of an animate model with an agency in social learning is well studied in primates; however, it is generally in conjunction with manual action tasks [62]. Our experiments, probably for the first time, highlights the importance of witnessing a model with an agency to learn the communicative act.

Another unexpected observation in this study was OT’s ‘failed’ (i.e., unvoiced) attempts to produce move-grunts, akin to the ‘silent’ vocalisations in the conditioning experiments with macaques [49,50]. During the 23 months of fieldwork on the same groups, we never observed anything similar in adults in any context (AD, personal obs). Previous attempts suggest it is difficult to train monkeys to vocalise on command [49,63]. In line with this, we noticed that OT also required significant physical efforts to elicit a successful call. Furthermore, we also observed that OT produced a few ‘wrong’ calls that resembled eagle alarm calls [64], despite the fact that there were no predators around and that OT did not show any anti-predatory behaviour. We speculate that OT was trying to produce a move-grunt but erroneously produced an alarm call.

These observations are also in line with recent findings on the neurology of primate call production. Coudé et al. identified vocalisation-selective neurons in the ventral premotor cortex, which fired only during the production of conditioned calls and remained inactive during silent vocalisations and spontaneous calls. Authors suggested that neural machinery required for voluntary vocalisation is not as fully formed as neural mechanisms involved in the spontaneous call in primates [49]. Notably, the volitional vocalisations also require greater efforts in motor preparation compared to volitional manual actions in primates [65]. In line with this, recent evidence suggests the correct response rate is much lower in vocal task compared to manual action task, probably due to separate neural networks for volitional vocalisations and manual actions in the ventrolateral prefrontal cortex [66]. This brain area is a homologue of the human Broca’s area known to play a crucial role in human speech. The duality in neural networks might also explain the higher success of social learning for manual tasks than our results [37,38].

Overall, our results and behavioural observations support the neurobiological evidence and suggest that non-human primates can probably socially learn the novel call usage from appropriate virtual stimuli. However, they have limited cognitive control over calls due to a lack of neural machinery, making it rather difficult to elicit a pre-existing call in an emotionally and functionally novel context. Contrary to our initial prediction, a possibility that move-grunts are not completely decoupled from the specific affective state could also explain the difficulty in its production in a novel context. In other words, we cannot discount the possibility that monkeys might have understood the task by watching the demonstration video; however, they were not equipped to perform it [67].

To our knowledge, this is the first study to use the social learning paradigm to test for functional flexibility of vocalisations in wild primates experimentally. We acknowledge that our sample size is small and lack rigorous statistical analyses. Results can be considered suggestive as only one subject seemed to learn the novel usage. However, it is comparable to other experiments conducted in captivity [22,24,68]. Furthermore, we have also provided comprehensive qualitative behavioural observations, which could be useful for future studies, and opened a new possible experimental paradigm that can be used on highly habituated and provisioned populations of different primate species. Future studies with an improved experimental design are required to confirm our results. In the future, researchers can use multiple remote-controlled food dispensers, which might reduce any effects arising from the experimenter’s proximity and improve the participation of subjects. A close-up camera and audio recorder that can record facial and mouth movements would help record behaviours like silent vocalisations and calls systematically. Acoustic analyses of the elicited calls were irrelevant for this study as rewards were based on the experimenter’s immediate classification of the call, and bad weather conditions did not yield good recordings. However, post-hoc acoustic analysis of the well-recorded volitional calls is critical to understanding the differences in calls elicited in natural and novel contexts [49].

Furthermore, we argue that experiments in captive settings that test for the cognitive control of vocalisation with neuronal recordings can adopt the social learning paradigm. It is ecologically more relevant than the previously used operant conditioning, and we speculate that it would provide substantial insights into the neural mechanism of volitional call production. Finally, similar experiments with anthropoid apes and prelinguistic children are necessary for the comparative analyses to understand the evolution of functionally flexible sounds in the primate lineage, offering fundamental insights into human language evolution.

## Author contribution

AD conceived the study with help from KZ. AD conducted the experiments analyses and wrote the first draft, which KZ reviewed. AD and KZ wrote the final version of the manuscript.

## Acknowledgements

AD would like to thank Maria Teresa Navarette and Mábia Biff Cera for suggestions on the experimental design of Experiment 1. Finally, we would like to thank all the volunteers, researchers, and managers at the Inkawu vervet project for their help during data collection.

