## Supplementary material for "Socially learned arbitrary call use in a wild primate"

### 1.1 Chapter 3: Appendix

Figure S1: Spectrogram (Hamming window at 1024 DFT and 93.8% overlap) of the naturally recorded move-grunt call used for the Experiments

Time (s)

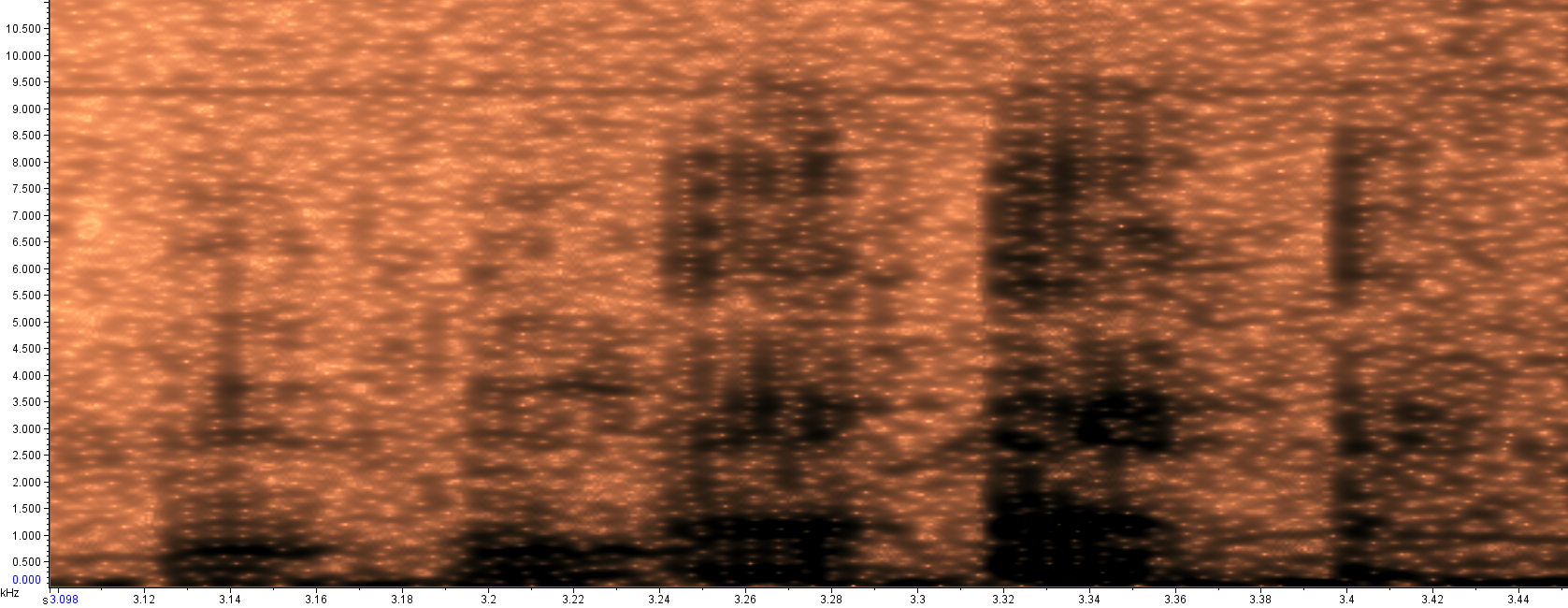

Frequency KHz

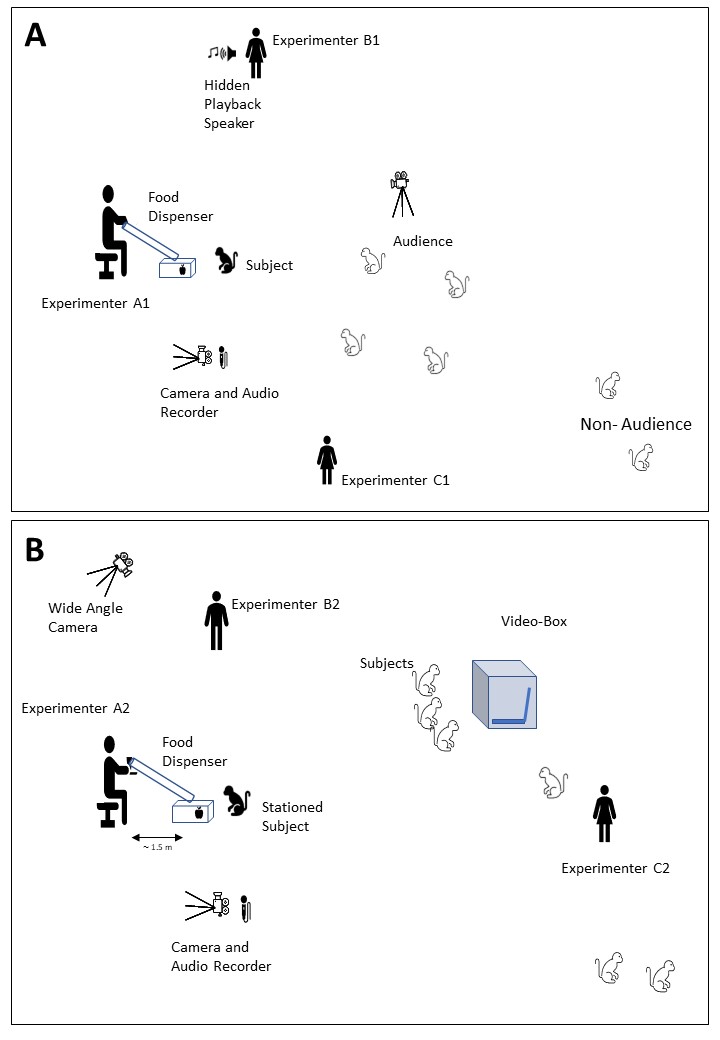

Figure S2: Heuristic illustrations (not to scale) of experimental setups (A) Experiment 1: Social learning of auditory demonstrations, (B) Experiment 2: Social learning of audio-visual demonstrations

Figure S3: Number of subjects participated per training day for Experiment 1: Social learning of auditory demonstrations

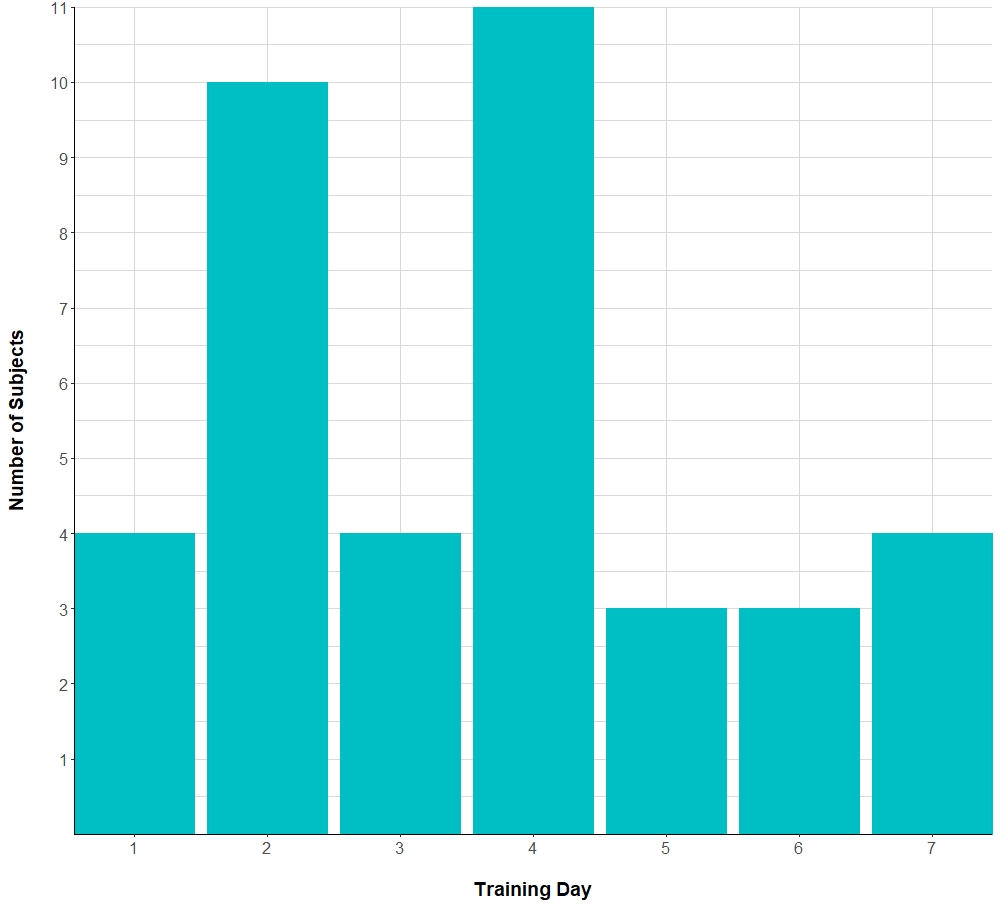

Figure S4: Subject wise participation in Experiment 1: Social learning of auditory demonstrations

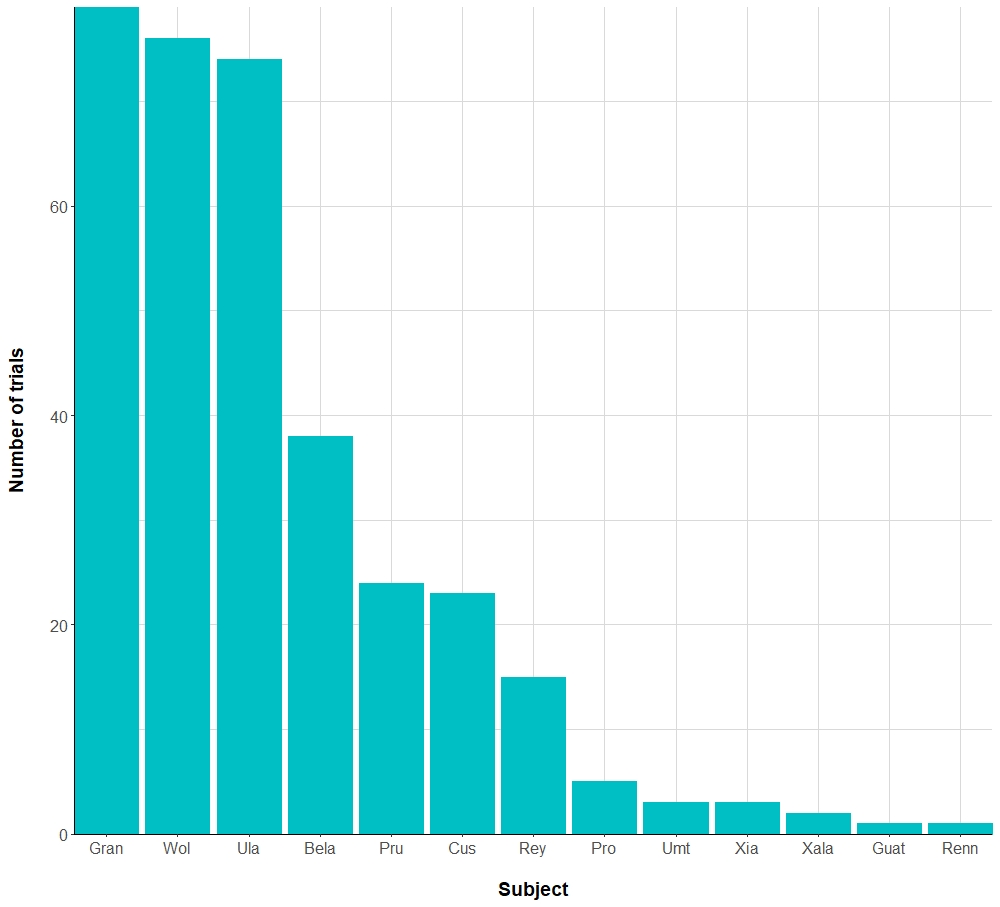

Figure S5: Focal behaviours by subject OT across sessions in Experiment 2: Social learning of audio-visual demonstrations

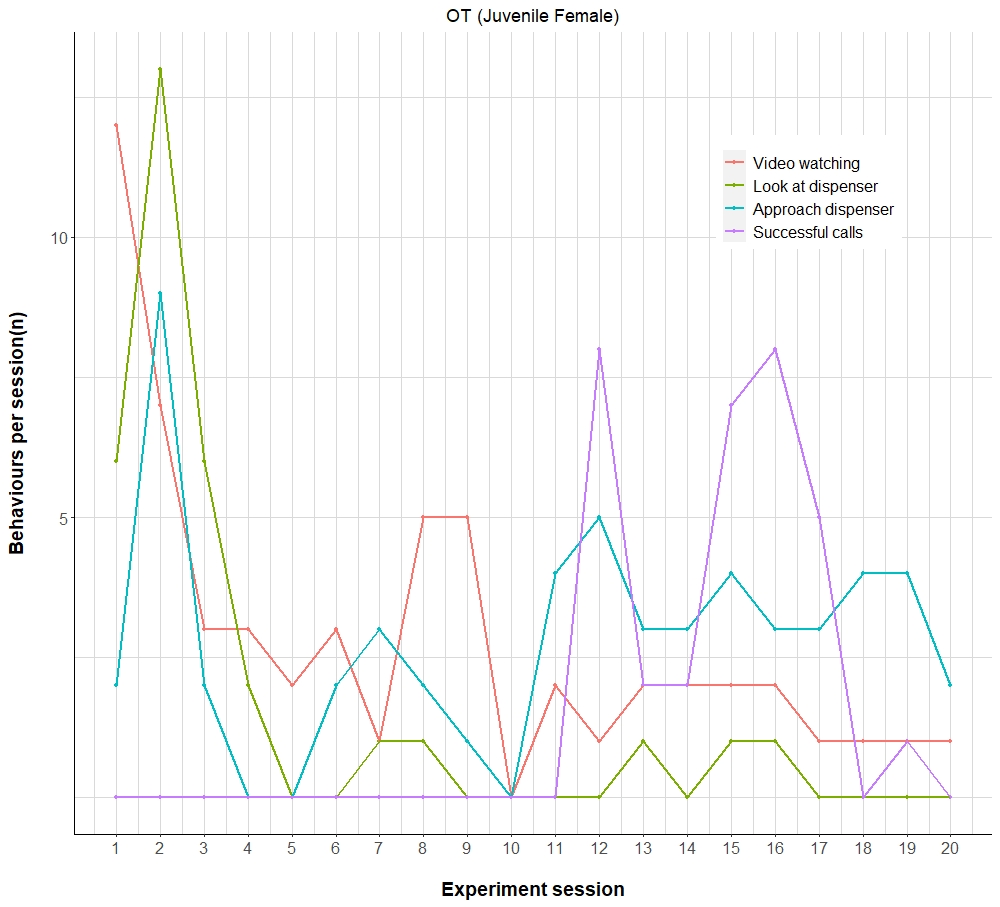
